# Amplified MS2 labeling reveals heterogeneous dynamics of RNA granules in the live murine brain

**DOI:** 10.64898/2026.02.03.703574

**Authors:** Hyungsik Lim, Grace P. Cooper

**Affiliations:** School of Optometry, Indiana University, Bloomington, IN 47405

**Keywords:** RNA imaging, MS2 labeling, hippocampus, two-photon microscopy

## Abstract

Dynamics of messenger ribonucleic acids (mRNAs) and the complexes with associated proteins (RNPs), also known as RNA granules, provide post-transcriptional control of gene expression. This regulation is crucial for cell-type diversity and activity-dependent plasticity of the mammalian central nervous system (CNS). However, elucidating the physiological significance of RNA granules has been hampered by the lack of technologies for probing them in live mouse brain. Here, we describe a novel method to visualize RNA granules in native CNS tissues in vivo. To amplify the fluorescence signal from single mRNAs incorporating MS2 stem loops, MS2 capsid protein (MCP) was conjugated with 4 tandem superfolder green fluorescent proteins (sfGFPs). Using MCP-4×sfGFP, a significant population of RNPs could be detected in specific cells of the CNS, enabling new findings, e.g., remarkable heterogeneity of *Actb* mRNA dynamics across neurons and glial cells. The highly sensitive in vivo RNA imaging could be useful for illuminating the regulation of the dynamics of RNA granules in naive and diseased animals.

**Significance:** RNA granules, complexes of RNAs and binding proteins, play a vital role in regulating gene expression and mRNA dynamics. Despite the importance, monitoring their behavior in functional neural tissue has proven technically challenging, in part due to optical turbidity and weak fluorescent tags. In this study, we present a new approach to better observe RNA granules in specific cells of murine brain through the use of amplified fluorescent signals and advanced imaging techniques. Our method provides a powerful means to analyze the properties of RNA granules within neurons and glial cells in vivo, offering valuable insights into healthy and neurodegenerative brains.

## 1. Introduction

Dynamics of ribonucleic acids (RNAs) are an element of post-transcriptional control of gene expression (1-4). After the synthesis and processing of messenger RNAs (mRNAs), various types of ribonucleoprotein (RNP) complexes are assembled in the cytoplasm, also known as RNA granules, which modulate the stability, trafficking, and translation of mRNAs (5-7). Many aspects of RNA granules are still unknown, particularly the physiological significance in vivo, although their dysregulation is implicated in neurodegenerative disorders (8-10). Transient RNA granules have been hypothesized to enable many polarized cells of the central nervous system (CNS), which is characterized by diversity of cell types and activity-dependent plasticity. To understand how dynamics of RNPs support these CNS characteristics and how failure gives rise to diseases, the molecules must be observed in their native environment where cellular identity and intrinsic physiology are preserved. The MS2 method allows nascent transcripts and endogenous mRNAs to be visualized in vivo by inserting multiple copies of stem loop of MS2 bacteriophage, i.e., MS2 binding sequence (MBS), into the gene of interest and expressing MS2 capsid protein (MCP) which targets the MBS, fused with green fluorescent protein (GFP) (11). Then, the RNA of interest binds numerous MCP-GFPs, amplifying fluorescent signals for detection. However, despite its widespread success in live-cell studies, the conventional MS2 method falls short of imaging single RNP particles in thick brain tissue due to turbidity degrading the resolution and signal-to-background ratio. Therefore, an advanced technique is required for elucidating the in vivo function of RNA granules in the context of pathophysiology of the mammalian brain.

Here we present a novel MS2 labeling technique that increases sensitivity for detecting endogenous mRNAs in live tissues. The fluorescence from individual mRNAs is increased by tagging MCP with 4 tandem GFPs, sufficient for visualizing single RNP particles in the turbid media. The amplified MS2 labeling is demonstrated for measuring the dynamics of *Actb* mRNAs in neuronal and glial cells of the murine brain.

## 2. Results

### Sensitive detection of RNA granules in fresh brain slices

To increase fluorescence per individual RNA molecule, we tagged MCP with four copies of GFP (Fig.1(a)). These tandem GFPs provide compounded amplification that is orthogonal to multiple MBS repeats. To minimize the risk of forming RNA aggregates, superfolder GFP (sfGFP) was used, i.e., a GFP variant with robust folding mechanics (12). For delivering MCP-4×sfGFP into live tissues, an adeno-associated virus (AAV) vector was designed with the human ubiquitin C (UbC) promoter and the woodchuck hepatitis posttranscriptional regulatory element (WPRE). The number of tandem GFPs was limited by the capacity of AAV packaging. The MCP-4×sfGFP sequence was approximately 3.3 kb and the total size of insert, including the inverted terminal repeats (ITRs), was ∼4.9 kb. To target *Actb* mRNAs, a knock-in mouse strain *Actb*^*MBS/MBS*^ was employed, containing 24 repeats of MBS in the 3’ untranslated regions (UTR) of both *Actb* alleles (13). As a result, each *Actb* mRNA could harbor up to 48 MCP-4×sfGFPs, or a total of 192 sfGFPs. After stereotaxic injection of high-titer AAVs, the acute brain slices of the *Actb*^*MBS/MBS*^ mouse were imaged by two-photon microscopy (TPM). In the hippocampi expressing MCP-4×sfGFP, bright fluorescent spots could be identified without additional image postprocessing (Fig.1(b)). By contrast, the *Actb*^*MBS/MBS*^ brain expressing MCP-1×GFP displayed far fewer punctate features above the background. To confirm the fluorescent puncta were *Actb* mRNAs, the images were compared to negative control, i.e., wild-type (WT) mice without MBS injected with MCP-4×sfGFP. Bright spots were absent in the WT brain (Fig.1(c)), suggesting that the signals were associated with MBS, thus likely from *Actb* mRNAs. The intensity of particles was heterogenous, possibly due to the variable number of mRNAs. Their sizes were close to the optical resolution of microscopy, but the size alone could not determine whether some particles contained multiple mRNAs. *Actb* RNA granules exhibited an uneven spatial distribution in the hippocampus, being much more enriched in the somata than the neuropils. Most RNA granules were extremely mobile, but a small number of them were immobilized near the perinuclear space (Fig.1(d), Movie S1). The movement of these RNA granules was confined primarily to the soma and also excluded from the intracellular organelles (Fig.1(d)). Notably, nuclear mRNAs were scarce and RNA granules in the dendrites were also relatively fewer than in the soma. The dendritic RNPs were much dimmer than somatic particles, suggesting fewer copies of *Actb* mRNAs.

**Figure 1.**
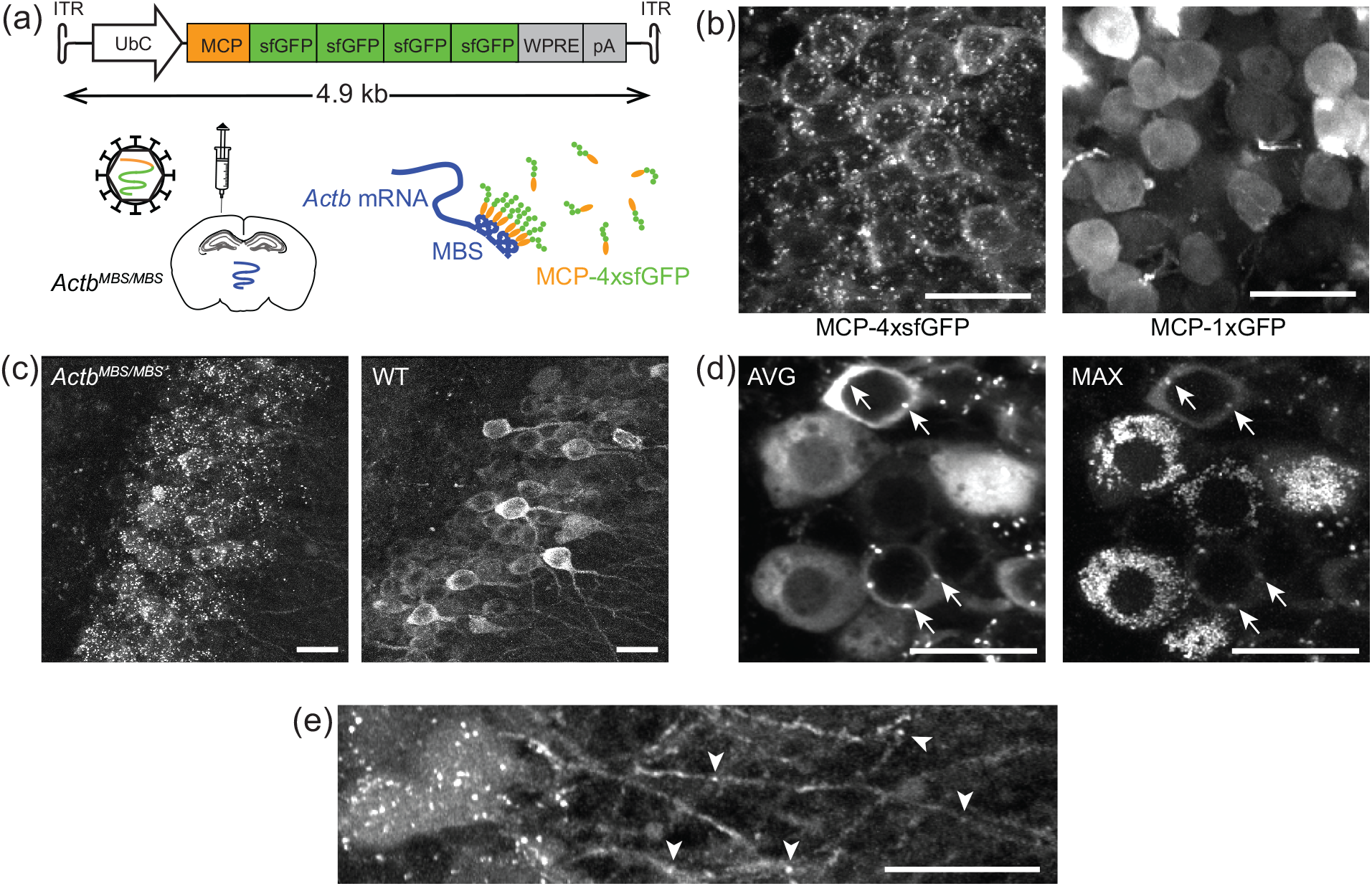
Sensitive detection of mRNAs with MCP-4×sfGFP. (a) The DNA construct and labeling procedure. pA, poly-adenylation signal. (b) The hippocampus of *Actb*^*MBS/MBS*^ mice expressing MCP-4×sfGFP vs. MCP-1×GFP. (c) *Actb*^*MBS/MBS*^ vs. WT mice expressing MCP-4×sfGFP. (d) Average and maximum projections of time-lapse images of *Actb*^*MBS/MBS*^ brain expressing MCP-4×sfGFP. Arrows, RNPs anchored near the perinuclear space. (e) RNPs in the dendrites (arrowheads). Scale bars, 20 µm.

### Cre-dependent MCP-4×sfGFP for imaging RNP particles in specific cells

Dynamics of mRNAs are highly variable depending on cell type, which is difficult to resolve with the UbC promoter driving the expression of MCP-4×sfGFP in multiple cell types within the brain. To isolate cell-type-specific dynamics of *Actb* RNPs, we engineered a Cre-dependent MCP-4×sfGFP by flanking it in a double-floxed inverted open (DIO) reading frame (14), i.e., DIO-MCP-4×sfGFP (Fig.2(a)). Consequently, while every *Actb* mRNA in the *Actb*^*MBS/MBS*^ mouse contains MBS, only those in *Cre+* cells were fluorescently labeled where DIO-MCP-4×sfGFP was inverted by Cre-lox recombination. A nuclear localization signal (NLS) was included to improve the contrast for cytoplasmic RNA granules by sequestering free MCPs to the nucleus (11). A truncated WPRE, i.e., W3 (15), was employed to keep the total size of transgene below the AAV packaging capacity. The properties of neuronal RNA granules have been extensively studied (16-18). To target them, *Actb*^*MBS/MBS*^ mice were crossed with *CaMKIIα-Cre* and *Grik4-Cre* strains (19, 20). Furthermore, for the visualization of RNA granules in glial cells, mice were used expressing Cre under the control of glial fibrillary acidic protein (Gfap), i.e., *Gfap-Cre* (21, 22). The hybrids that were homozygous in MBS and positive in Cre, referred to as *Actb*^*MBS/MBS*^*;CaMKIIα-Cre, etc* (Fig.2(b)), received AAV injection. Fig.2(c) depicts *Actb* RNPs in the *Cre+* cells within the CA3 region. RNPs exhibited significant enrichments in the neuronal somata (Fig.2(c), (d)), like Cre-independent MCP-4×sfGFP. In *Actb*^*MBS/MBS*^*;Gfap-Cre* mice, however, it was difficult to delineate RNA granules specific to glial cells due to substantial Cre expression in nearby pyramidal neurons (Fig.2(e)).

**Figure 2.**
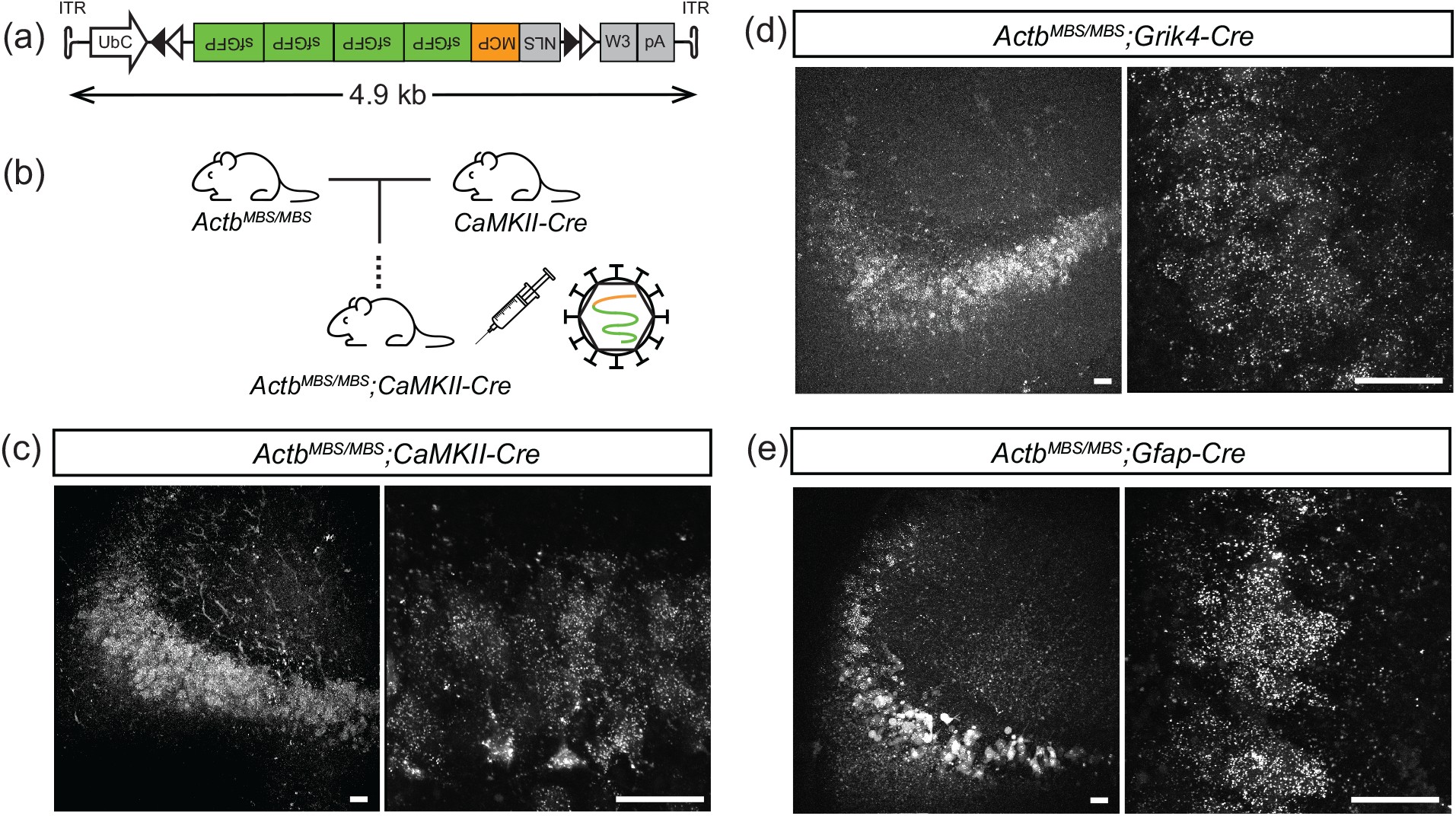
Visualizing *Actb* mRNAs in specific cells. (a) The DNA construct for Cre-dependent MCP-4×sfGFP. (b) The creation of hybrid *Actb*^*MBS/MBS*^ Cre mouse strains. (c-e) *Actb* mRNAs in the *CaMKIIα-Cre+, Grik4-Cre+*, and *Gfap-Cre+* cells of the hippocampus. Scale bars, 40 µm.

### Neuronal *Actb* RNA granules were heterogenous in the hippocampus

With the ability to visualize *Actb* mRNAs in specific cells, their properties could be measured free from the heterogeneity across cell types. First, we determined the gain by which the fluorescence of mRNAs was amplified, i.e., whether MCP-4×sfGFP yielded 4 times larger signals compared to MCP-1×GFP. AAVs carrying DIO-MCP-4×sfGFP or DIO-MCP-1×GFP were injected, acute brain slices were imaged by TPM. Representative images of the hippocampi of *Actb*^*MBS/MBS*^*;CaMKIIα-Cre* are shown in Fig.3(a). MCP-4×sfGFP allowed nearly thirteen times more RNA particles to be detected compared to the standard MCP-1×GFP. For precise quantification of the image contrasts, RNA particles and the soma background were segmented (Fig.3(b)). The intensity of RNA particles labeled by MCP-4×sfGFP, relative to the somatic background, was on average 2.75 times higher than those labeled by MCP-1×GFP (Fig.3(c)). The less than anticipated amplification factor can be explained by a few reasons. First, it is possible that the somatic background was higher with MCP-4×sfGFP. Second, and more importantly, the composition of the labeled RNA particles could be different between MCP-4×sfGFP and MCP-1×fGFP. Intensity histograms showed multiple subpopulations of RNA particles visualized with MCP-4×sfGFP (Fig.3(d)), indicating neuronal RNA granules were heterogenous. The five most significant components, out of 9 yielded by the maximum likelihood Gaussian mixture model with the lowest Akaike information criteria (AIC) value, displayed quantal intensities with mode spacing comparable to the intensity of background. Assuming the uniform increment of intensity was due to the integer number of *Actb* mRNAs in the RNPs, it can be concluded that particles with more than 2 mRNAs were detectable with MCP-4×sfGFP. In contrast, the histogram of RNA particles labeled by MCP-1×GFP did not show discernible subpopulations, most likely because only a subset of RNPs containing many *Actb* mRNA molecules could be visualized. The skewed sampling by MCP-1×GFP that favored larger RNPs likely led to a lower than predicted gain. Overall, quantitative analysis validated the sensitivity of MCP-4×sfGFP for visualizing a comprehensive set of RNA granules.

**Figure 3.**
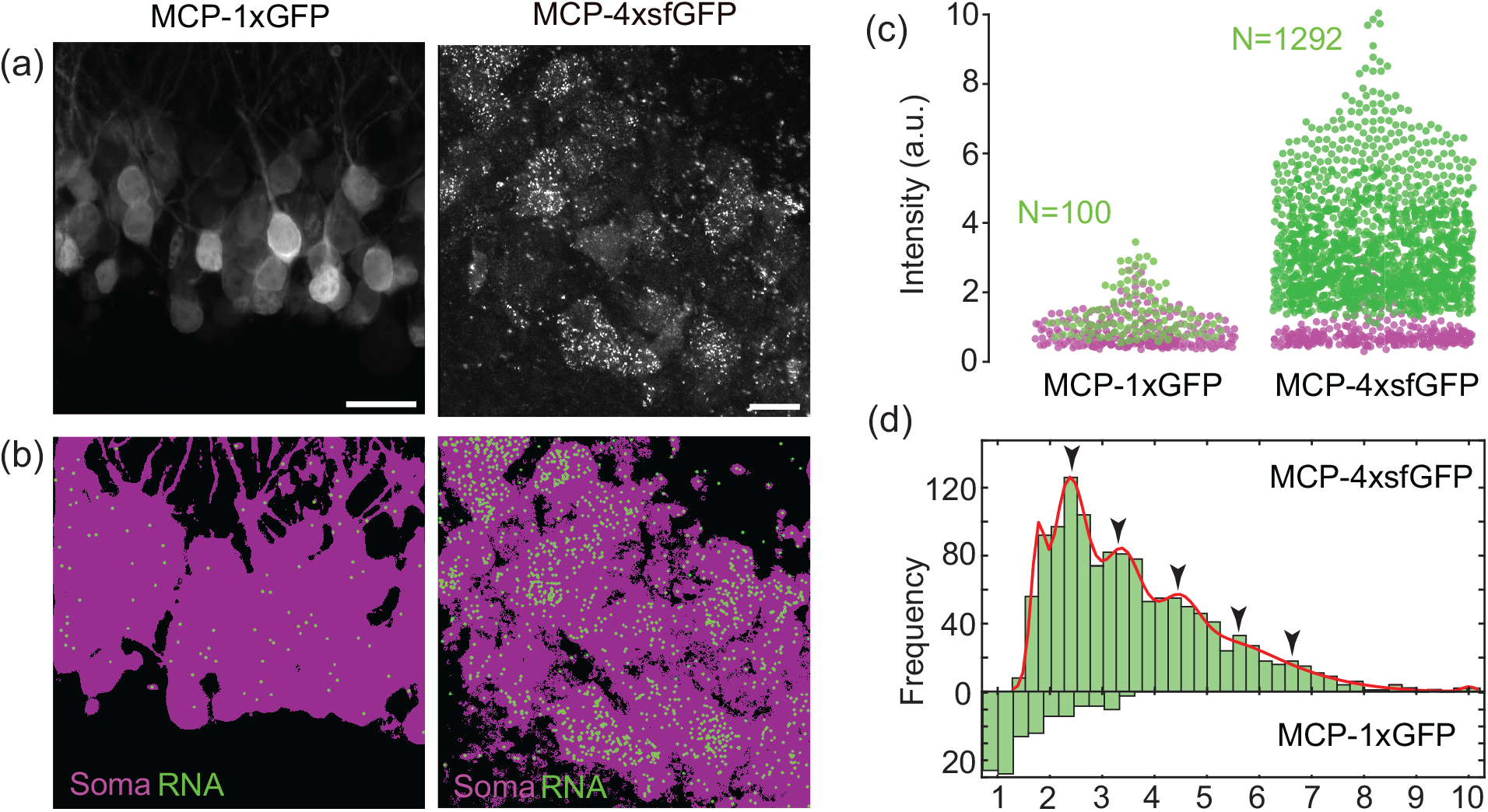
Quantitative analysis of *Actb* mRNAs in neuronal RNA granules. (a)*Actb*^*MBS/MBS*^*;CaMKIIα-Cre* brains expressing MCP-1×GFP vs. MCP-4×sfGFP. Scale bars, 20 µm.Segmentation of RNA particles (green) and somatic background (magenta). (c) The intensity distribution of RNA particles vs. background. (d) Histogram of the normalized intensity of detected RNA particles. Red line, the maximum likelihood Gaussian mixture model. Arrowheads, 5 major components of the largest proportions with modes at 2.34, 3.36, 4.47, 5.21, and 6.42.

### Movements of neuronal RNA granules were mostly stochastic and independent of active transport

Next, movement of neuronal RNA granules was analyzed. In hippocampal *CaMKIIα+* and *Grik4+* neurons, *Actb* mRNAs were largely somatic and exhibited random motion (Fig.4(a), Movie S2). The entire cytoplasm minus organelles was accessible to mobile RNA granules (Fig.4(b)). We asked whether the diffusive movement of *Actb* RNA granules depended on the cytoskeleton or molecular motors. To address this, pharmacological agents were employed, including Jasplakinolide, an actin stabilizer, and nocodazole, a microtubule depolymerizer. Tetracaine was also used as a promiscuous inhibitor of kinesin motor proteins (23). We found that the diffusive motion of *Actb* mRNAs did not respond to the 1-hour treatment with 10-µM Jasplakinolide or 33-µM nocodazole (Fig.4(c)). Their speed or accessibility were unaffected qualitatively, indicating that the diffusive motion did not require dynamic actin filaments or the integrity of microtubules. Similarly, intervention with 5-mM tetracaine for >1 hour did not alter the overall dynamics. The results suggested that diffusive RNA granules were not engaged with the cytoskeleton or motor proteins. Single-particle tracking (SPT) was performed to characterize the diffusive movement quantitatively (Fig.4(d)). It is common to examine random motions via the mean square displacement. However, due to short track lengths arising from the 2D image plane sampling the 3D movement of RNP particles, it was more appropriate to evaluate the jump distance distribution, which can resolve distinct mobile populations undergoing normal diffusion (24, 25). The best fit to a 2D diffusion model yielded two distinct species with diffusion constants of 0.026 and 0.68 µm^2^/sec with fractions of 21 and 79%, respectively (Fig.4(e)). The diffusion constant of mobile species was consistent with the range determined previously using live *Actb*^*MBS/MBS*^ cells (26). Thus, we conclude that the diffusive movement of *Actb* RNPs were not altered significantly by binding with multiple copies of MCP-4xsfGFP. Our observations that neuronal RNA granules were mostly stochastic are in contrast to the previous studies showing directional motion and localization of *Actb* mRNAs are mediated by active transport (27, 28). The difficulty in detecting directional RNA granules could arise from technical reasons: Our frame rate,approximately 1 Hz of typical scanning beam microscopy, was much slower than the previous studies employing wide-field illumination so the temporal resolution may not be adequate to capture brief transport on shorter time scales. Furthermore, unlike in live-cell imaging where the full neurite lies within a field of view, the spatial range for tracing directional RNA granules was limited due to the 2D image plane intersecting the 3D tissue. We also speculate that the fraction of mRNAs in directional motions might be inherently low at any time point, which is discussed in the next section.

**Figure 4.**
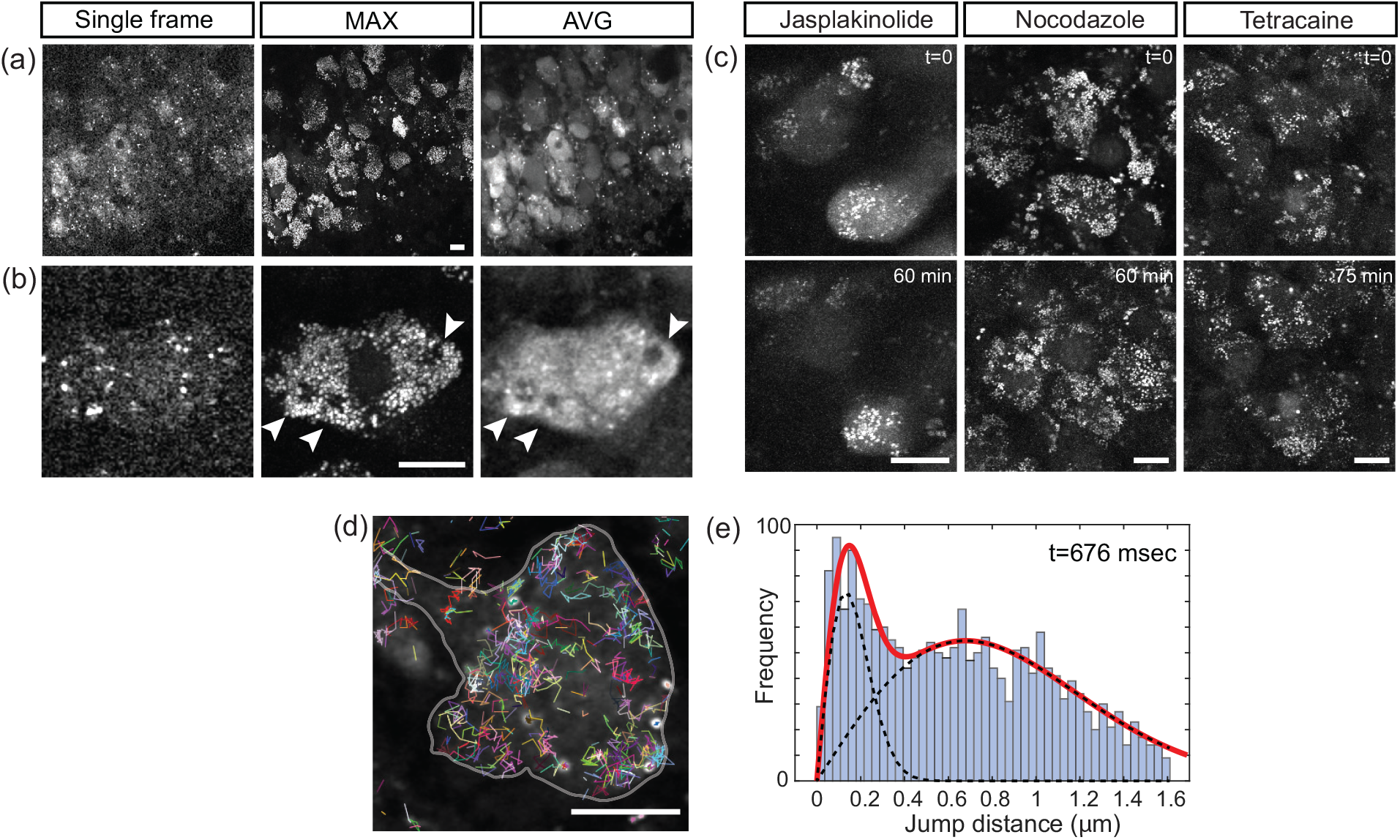
Movements of neuronal RNA granules. (a), (b) Single frame, maximum and average projections of time-lapse images of MCP-4×sfGFP expressing cells in the *Actb*^*MBS/MBS*^*;Grik4-Cre* hippocampus (time=162 sec). Arrowheads, cytoplasmic compartments limiting the access of RNPs. (c) Max projections of time-lapse images before and after the treatments of Jasplakinolide (10 µM), nocodazole (33 µM), and tetracaine (5 mM). (d) Single particle tracking traces overlaid with average projection. (time=33.8 sec). (e) Jump distance distribution with the best fit to two-component 2D diffusion model (red). Dashed lines, each component. Scale bars, 10 µm.

### Glial *Actb* RNA granules exhibited distinguishable dynamics

Localization and local translation of *Actb* mRNAs have been studied using cultured neurons and fibroblasts (29, 30). Evidence suggests mRNAs localize also in other polarized cells of the brain, e.g., radial glia (31), oligodendrocytes (32), and astrocytes (33, 34). However, the in vitro cultures may not recapitulate the actual physiology of the CNS, thus unideal for interrogating mRNA dynamics. We employed MCP-4×sfGFP for visualizing *Actb* mRNAs in non-neuronal cells of the brain. As an alternative to *Actb*^*MBS/MBS*^*;Gfap-Cre*, which expressed Cre in neurons as well (Fig.2(e)), we employed a synthetic Gfap promoter, i.e., GfaABC1D (35). The AAV for delivering GfaABC1D-MCP-4×sfGFP (without an NLS) was produced (Fig.5(a)). After AAV injection, the acute brain slices were imaged. RNA granules in *Gfap+* cells of the hippocampus were visualized as punctate dots and their identity as *Actb* RNPs was confirmed by negative control with WT (Fig.5(b)). Interestingly, *Actb* RNPs exhibited dynamic properties that were visibly distinguishable from those in neurons. Whereas neuronal RNPs were mostly enriched in the cytoplasm, in glial cells a substantial fraction of them were localized to the distal processes. Also, regardless of localization, i.e., in the soma or processes, *Actb* RNA granules in *Gfap+* cells were much less mobile (Movie S3). *Gfap+* cells are not homogenous, encompassing multiple cell types across the brain. To dissect mRNA dynamics in a region-specific manner, two characteristic *Gfap+* cells were examined in the native niche, e.g., mature astrocytes in the CA1 and radial glia-like cells in the subgranular zone (SGZ) of the dentate gyrus (DG). The latter is adult neural stem cells so we anticipated the dynamics of RNPs therein might emulate that of developing neurons (29) rather than terminally differentiated neurons. Both astrocytes and radial glia-like cells exhibited substantial localization of *Actb* RNPs (Fig.5(c)-(f)). In the radial glia-like cells, modulated intensities within the processes suggested relatively small copy numbers of *Actb* mRNAs (Fig.5(e), (f)). The distinct mRNA dynamics observed in *Gfap+* cells could derive from diversified function of *Actb* mRNAs.

**Figure 5.**
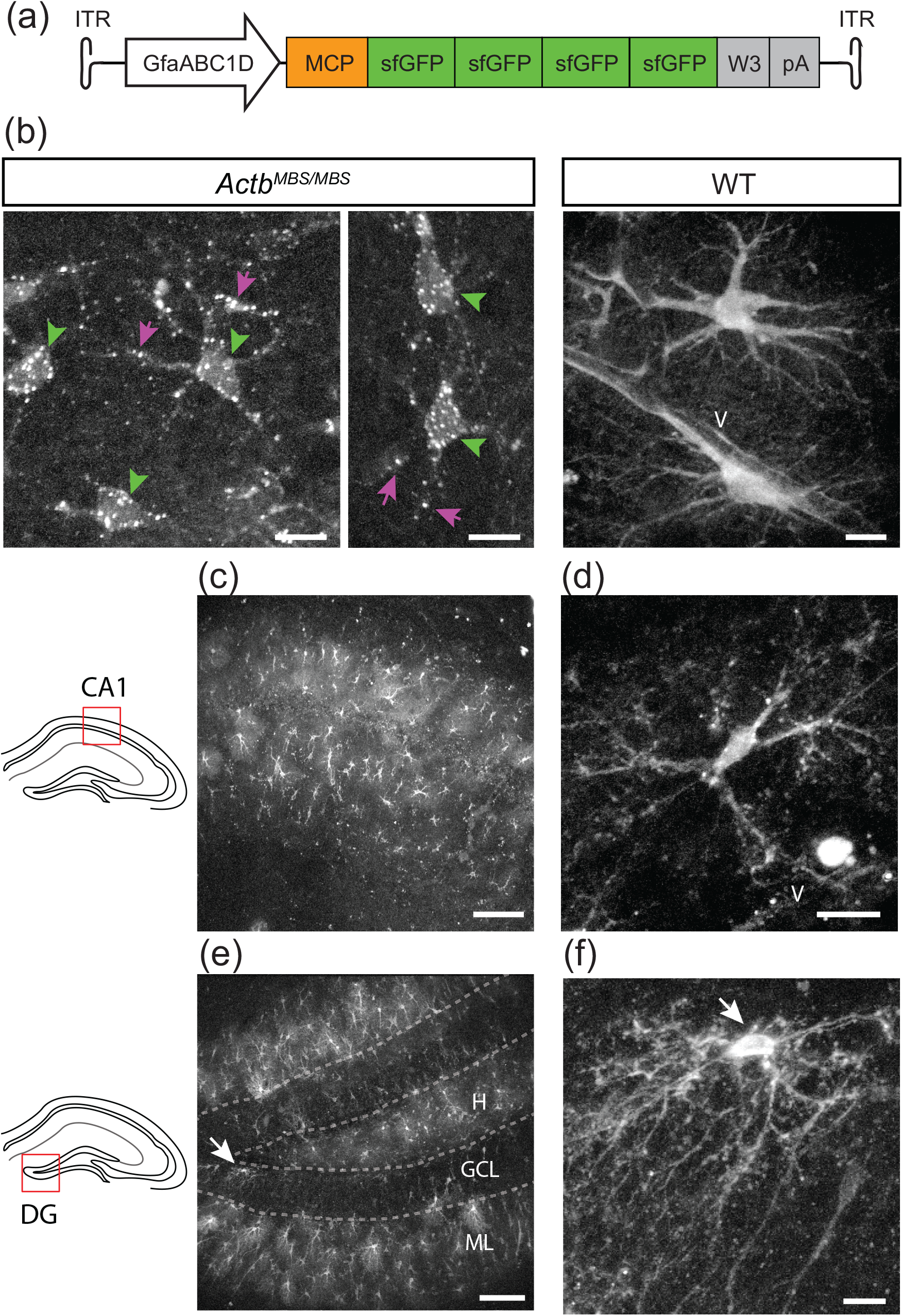
Visualizing *Actb* RNPs in the hippocampal glial cells. (a) The DNA construct for expressing MCP-4×sfGFP under the control of GfaABC1D promoter. (b) *Actb*^*MBS/MBS*^ vs. WT mice expressing GfaABC1D-MCP-4×sfGFP. *Actb* RNPs in the soma and processes (arrowheads and arrows, respectively). V, blood vessel. (c) MCP-4×sfGFP in *Gfap*+ cells of the CA1 region. (d) Mature astrocyte in CA1. (e) MCP-4×sfGFP in *Gfap*+ cells of the DG region. H, hilus; GCL, granular cell layer; ML, molecular layer. (f) Radial glia-like cell in the SGZ (arrow). All intensities are shown on the log scales. Scale bars: 100 µm ((c), (e)) and 10 µm (all others).

## 3. Discussion

We have presented a new MS2 labeling strategy for upgrading fluorescence from single RNA molecules. With enhanced MCP signals, a wider range of RNPs could be detected in the murine brain than previously possible, which is a significant advance over live-tissue RNA imaging based on MCP-GFP (36, 37). The sensitivity of MCP-4×sfGFP nearly matches that of in vitro RNA labeling, such as fluorescence in situ hybridization (FISH), revealing a comprehensive set of RNPs. Furthermore, we have resolved RNAs in specific cells, as suitable for unraveling the cell-type-dependent dynamics, which is difficult to achieve with FISH. Our method is comparable to another recent technique of amplifying MS2, in which each mRNA can be labeled with up to 384 GFPs for 8 MBS repeats (38). Utilizing the Suntag system (39), the method conjugates multiple GFPs per MCP protein via two separate modules, i.e., MCP-24×Suntag and single-chain variable fragment antibody tagged with GFP (scFv-GFP). As a result, attaining the predicted contrast hinges on balancing the bipartite stoichiometry, i.e., 24 scFv-GFP per MCP. MCP-4×sfGFP does not have such requirements.

Our method could be improved by considering a few issues. MCP-4×sfGFP preferentially visualizes a subset of RNA granules with many copies of mRNAs, omitting those with one or two mRNAs. The latter includes an important class of functional RNA particles, e.g., in neurites (40). To capture the whole population of RNPs, MCP signals could be increased further with more GFPs in tandem, which is feasible by employing lentiviral vectors with a larger packaging capacity (8-12 kb). However, there are potential caveats: A reporter protein larger than MCP-4×sfGFP (∼125 kDa) may be difficult to pass through the nuclear pore (41), compromising the efficiency of targeting nuclear RNPs. Also, binding with numerous bulky MCP proteins could hinder or alter the movement of mRNAs. Lastly, unusually large reporter proteins could increase the potential for non-physiological artifacts, such as RNA aggregates (42).

With MCP-4xsfGFP and live tissue imaging, we have acquired new findings regarding dynamics of *Actb* RNA granules, e.g., surprising diversity of mRNA dynamics across cell types for this ubiquitous housekeeping gene. Our data also confirms the prior knowledge from FISH or live-cell imaging of primary or cultured cells, e.g., somatic neuronal RNA granules display random diffusive motions that are independent of the cytoskeleton or motor proteins. However, MCP-4xsfGFP has yielded some observations that deviate from the previously documented behaviors (28, 43-45). For instance, we have found *Actb* RNPs are relatively scarce in the neuropils of the hippocampus and there are few motor-driven transports or directional *Actb* mRNAs. In addition to the technical difference, it could be also due to the low frequency of long-range active transport, which hasn’t been quantified systematically. In a steady state of differentiated neurons in vivo, a nonnegligible fraction of directional RNPs would lead to an imbalance in the intracellular distribution. An equilibrium with substantial bidirectional active transport might be less favorable energetically to an ensemble of stochastic RNA granules. Conceivably, cytoplasmic RNA granules may undertake intracellular surveillance until they receive appropriate cues for anchoring/active transport. The transition between different classes of motion could thereby provide a layer of regulation in gene expression. RNA imaging with MCP-4xsfGFP is an ideal tool to image transient neuronal granules in live tissues of various animal models.

Also, we have found the distribution and dynamics of *Actb* RNA granules in the glial cells are remarkably heterogeneous and distinguished from those in the neurons. Substantial localization of *Actb* RNA granules in the distal processes of *Gfap+* cells, including astrocytes and radial glia-like cells, may stem from a greater need for the local synthesis of β-actin protein. The *cis*-element and *trans*-acting factors underlying *Actb* mRNA localization have been identified using in vitro fibroblasts and neurons (46-48). The same mechanism is likely to be involved in the localization of *Actb* mRNAs in *Gfap+* cells. However, there are open questions, e.g., which specific molecules facilitate the cell-type variability. It seems feasible that MCP-4×sfGFP can unravel the cell-specific regulation in the native niche of brain circuits. *Gfap+* glia includes astrocytes, a major cell type accounting for approximately 30% of the cells in the mammalian CNS with many diverse functions in synaptic formation and plasticity, regulation of blood flow, and metabolic support of neighboring neurons. Direct observations of dynamic RNA granules of astrocytes in live brain tissue could illuminate the function in neuro-glia interactions, such as reactive astrogliosis and neuroinflammation and offer novel perspectives on their roles in age-related and neurodegenerative disorders.

## 4. Methods

### Animals

All procedures were approved by the Indiana University Institutional Animal Care and Use Committee (IACUC). All Cre strains and WT mice were obtained from Jackson Laboratory: B6.Cg-Tg(Camk2a-cre)T29-1Stl/J (#005359), C57BL/6-Tg(Grik4-cre)G32-4Stl/J (#006474),B6.Cg-Tg(Gfap-cre)73.12Mvs/J (#012886), and C57BL/6J (#000664). Knock-in mouse *Actb*^*MBS/MBS*^ was a gift from R.H. Singer. To create hybrid *Cre+ Actb*^*MBS/MBS*^ mice, *Actb*^*MBS/MBS*^ was bred with Cre mice and then the offspring animals of *Actb*^*MBS/+*^*;Cre+* were bred with *Actb*^*MBS/MBS*^. *Actb*^*MBS/MBS*^*;Cre+* mice were identified in the subsequent generations. Both male and female mice in the age range of 2 to 5 months old were used for experiments.

### Cloning MCP constructs

A plasmid for Cre-dependent MCP-1×GFP (pAAV-Ef1a-DIO-MCP-1×GFP-WPRE-hGHpA) was cloned by inserting MCP into pAAV-Ef1a-DIO-EGFP-WPRE-pA (Addgene plasmid #37084) using HiFi DNA Assembly Master Mix (New England Biolabs, E2621S). A plasmid for Cre-independent MCP-1×GFP (pAAV-Ef1a-MCP-1×GFP-WPRE-hGHpA) was produced by inverting pAAV-Ef1a-DIO-MCP-1×GFP-WPRE-hGHpA with Cre Recombinase (New England Biolabs, M0298S). For creating plasmids for Cre-independent and -dependent MCP-4×sfGFP (pAAV-UbC-MCP-4xsfGFP-WPRE and pAAV-UbC-DIO-SV40NLS-MCP-4xsfGFP-W3, respectively), DNA fragments were synthesized (Genewiz) and concatenated into a pAAV backbone containing a bovine growth hormone poly(A) signal (bGHpA). To produce a plasmid for expressing MCP-4×sfGFP under the control of GfaABC1D (pAAV-GfapABC1D-MCP-4×sfGFP-W3-bGHpA), a fragment including the promoter was amplified by polymerase chain reaction (PCR) from pAAV.GfaABC1D.PI.Lck-GFP.SV40 (Addgene plasmid #105598) and, along with MCP-4xsfGFP, ligated into a pAAV backbone.

### AAV

AAV packaging was performed by Charles River Laboratory, Franklin Biolabs, and Vector Biolabs. Capsid serotypes of AAV2, AAV5, or AAV9 were used. Briefly, HEK293 cells were co-transfected with ultrapure plasmids and recombinant AAV particles were purified by ultrahigh speed gradient centrifugation. Genome titer was determined either by droplet digital PCR (ddPCR) or by SYBR green qPCR. All AAVs were stored in -80°C until usage. The properties of AAVs are listed in Table S1.

### Stereotaxic injection

The AAV injection was performed on the stereotaxic frame (Kopf). The coordinates for the hippocampus were AP 2.0mm, ML 2.0mm, DV 2.0mm; for the cerebral cortex AP 2.0mm, ML 2.0mm, DV 2.0mm. The animal was anesthetized with 2% isoflurane, and the hair was shaved. The skin was sterilized, and a midline incision was made. The skull was drilled at the target position with a microdrill (Foredom). Using a 33-gauge needle and a Hamilton syringe, 1-µL AAV was slowly injected with at a rate of 0.1 µL/minute. After injection, the incision was closed with surgical staples.

### Acute brain slice preparation

Sectioning of fresh mouse brain was done using a vibratome (Leica) in an ice-cold artificial cerebrospinal fluid (ACSF) perfused with 95%O_2_/5%CO_2_. The thickness of slice was 250 µM. Immediately after sectioning, the brain slices were transferred to a bath containing ACSF perfused with 95%O_2_/5%CO_2_ and incubated at 37ºC for at least 1 hour for recovery.

### Pharmacology

Jasplakinolide and nocodazole (Santa Cruz Biotechnology, sc-202191 and sc-3518B, respectively) were dissolved in DMSO and added to the ACSF. Tetracaine hydrochloride (Sigma) was dissolved in water and added to the ACSF.

### Two-photon fluorescence microscopy

Two-photon microscopy was performed as described previously (36, 37). Briefly, a laser beam at 920 nm from a Ti:Sapphire laser (Chameleon Ultra, Coherent, Inc.) was used to excite GFP. The pulse duration was approximately 100 fs and the repetition rate was 80 MHz. A water-immersion microscope objective lens (HC FLUOTAR L 25x 0.95NA, Leica) was used to focus the excitation beam onto the sample. The average power at the sample was approximately 20 mW. Two detection channels employed narrow-bandpass filters (<20-nm bandwidth) and photomultiplier tubes (PMTs; H7422-40, Hamamatsu, Inc.). Images of 512×512 pixels were acquired at a frame rate of 0.74, 1.48, or 2.96 Hz.

### Image processing and data analysis

Image processing was performed using ImageJ (49). Image segmentation was obtained with an ImageJ plugin, Trainable Weka Segmentation (50). SPT was carried out with another ImageJ plugin, TrackMate (51). Data analysis was performed with MATLAB (MathWorks, Inc.). For the jump distance distribution, the equation for the probability density function *p*(*r, t*)of the two-component 2D diffusion model was 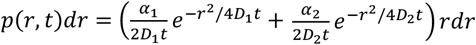, where r is the distance, t is the time, α_1_ and α_2_ are the fractions of two components, and *D*_1_ and *D*_2_are the diffusion constants.

## Acknowledgements

This work was supported by funding from the National Institute of Health (GM140841).

## Author contributions

H.L. designed research; G.P.C. and H.L. performed research; H.L. analyzed data; and H.L. and G.P.C. wrote the paper.

## Supporting Information for

### Tables

**Table S1.** The properties of AAVs.

|  | Vector | Capsid | Titer (GC/mL) | Source |
| --- | --- | --- | --- | --- |
| 1 | Ef1a- MCP-1×GFP | AAV9 | 1.44E13 | Charles River Laboratories |
| 2 | Ef1a-DIO-MCP-1×GFP | AAV2 | 7.68E12 | Charles River Laboratories |
| 3 | UbC-MCP-4xsfGFP | AAV5 | 8.76E12 | Franklin Biolabs |
| 4 | UbC-DIO-SV40NLS-MCP-4xsfGFP | AAV5 | 1.17E13 | Vector Biolabs |
| 5 | GfapABC1D-MCP-4×sfGFP | AAV5 | 4.67E12 | Franklin Biolabs |

**Movie S1 (separate file)**. Actb^MBS/MBS^ brain expressing MCP-4×sfGFP. 30× speed and time = 405 seconds.

**Movie S2 (separate file)**. MCP-4×sfGFP expressing cells in the *Actb*^*MBS/MBS*^*;Grik4-Cre* hippocampus. 30× speed and time = 162 seconds.

**Movie S3 (separate file)**. MCP-4×sfGFP expressing *Gfap*+ cell in the *Actb*^*MBS/MBS*^ hippocampus. 30× speed and time = 33.8 seconds.

